# Ancestry Influences on the Molecular Presentation of Tumours

**DOI:** 10.1101/2020.08.02.233528

**Authors:** Constance H. Li, Syed Haider, Paul C. Boutros

## Abstract

Epidemiological studies have identified innumerable ways in which cancer presentation and behaviour is associated with patient ancestry. The molecular bases for these relationships remain largely unknown. We analyzed ancestry associations in the somatic mutational landscape of 12,774 tumours across 33 tumour-types, including 2,562 with whole-genome sequencing. Ancestry influences both the number of mutations in a tumour and the evolutionary timing of when they occur. Specific mutational signatures are associated with ancestry, reflecting potential differences in exogenous and endogenous oncogenic processes. A subset of known cancer driver genes was mutated in ancestry-associated patterns, with transcriptomic consequences. Cancer genome sequencing data is not well-balanced in epidemiologic factors; these data suggest ancestry strongly shapes the somatic mutational landscape of cancer, with potential functional implications.

## Introduction

Racial differences in cancer are pervasive across myriad measures of cancer burden. Epidemiological studies have reported race-associated differences in incidence^1–4^, survival^3,5,6^, and mortality^1,2,7,8^ rates, amongst others. These persist despite advancements in cancer detection and treatment^9^. Across all cancer types, incidence rates in Black and White populations are comparable, but Black mortality rates are ~13% higher^1^. In contrast, all cancer incidence and mortality rates are lower in US Asian, Native Hawaiian and Pacific Islanders in the US^4^, and in UK East Asians and South Asians^3^ when compared with Whites. North American indigenous populations such as American Indians and Canadian First Nations have a risk of cancer death significantly higher than that for Whites, despite regional variation^1,10,11^.

There are more striking differences in the rates of diagnosis and mortality for specific tumour-types. Black men are twice as likely to be diagnosed with prostate cancer than White men, and twice as likely to die of their disease^1,2,12^. While breast cancer incidence rates in Black and White women have converged and are now comparable^1,2^, Black women still experience higher breast cancer mortality due in part to higher rates of aggressive triple negative disease and late stage at diagnosis^13–15^. In East Asians, the incidence rate of liver cancers is over twice that in Whites, and nasopharyngeal cancer incidence is six times greater: East Asians also have higher mortality for liver, nasopharyngeal, and stomach cancers^4,16–18^.

The causes of racial difference in cancer are multifactorial. Some differences in cancer survival are associated with differences in treatment effectiveness. For example, US studies have found that liver cancer survival in Black populations is lower after surgical interventions including hepatectomies and liver transplantation^19,20^. Black American men have been found to have poorer recurrence-free survival after radical prostatectomy^21,22^, but these differences in treatment response are at least in part, and perhaps mostly, attributable to differences in clinical and pathological characteristics at diagnosis and to socioeconomic factors^23,24^. Indeed, socioeconomic factors play an important role in cancer rates and outcomes: socioeconomic status directly affects critical variables such as living conditions and access to healthcare, and is strongly associated with health outcome throughout the world^25^. In other cases, comparing individuals across continents also confound life-style differences such as diet and environmental exposures, such as the prevalence of specific viruses. To better understand the causes of these differences in cancer incidence and mortality, many interacting and interrelated factors and concepts must be considered.

The concept of *ancestry* is itself closely related the concepts of *race* and *ethnicity*. Where *race* refers to groups distinguished by physical differences, *ethnicity* reflects differences by biological factors in addition to geographical, historical, belief, cultural and other factors. *Ancestry* as used here refers to genetic ancestry, or the line of descent of an individual’s genetic material. Genetic ancestry is directly imputed based on DNA sequencing^26^. Race, ethnicity, and ancestry are each associated with cancer burden. For instance, germline risk variants detected at different frequencies in different populations have been associated with differences in cancer risk^27^. Other population-specific risk variants have been described for breast^28,29^, prostate^30,31^, and lung^32,33^ cancer, and many are still of unknown significance. Our work focuses on describing the ancestry-associations of somatic genomic changes in cancer. However, it is important to note that studies of genetic ancestry cannot generally fully disentangle differences associated with genetics from differences associated with socioeconomic or other cultural and societal differences amongst populations.

We sought to understand how a tumour’s molecular profiles reflects the oncogenic processes and mutagenic exposures experienced by each unique patient. We used genetically imputed ancestry along with available (but inherently limited) lifestyle and clinical annotation data to model somatic features and identify features significantly associated with ancestry. Previous work associating somatic genomic changes with race, ethnicity or ancestry suggest differences in overall mutation burden^34,35^ in specific tumour-types like breast^36^, prostate^37^ and lung^38^ cancers, amongst others^39^. Studies of TCGA pan-cancer data have estimated genetic ancestry using SNP genotyping and compared African American-derived with European American-derived somatic alterations^40^, and examined the mRNA and methylation differences between ancestries^41^.

We add to this growing body of analyses examining ancestry-associated differences in cancer genomics by performing a pan-cancer, genome-wide study of ancestry-associated molecular differences, leveraging all available ancestry groups including those of European, East Asian, African, Admixed American and South Asian ancestry. Our comprehensive pan-cancer analysis leveraged 10,218 tumours of 23 tumour-types from The Cancer Genome Atlas (TCGA)^42^ and 2,562 tumours of 30 tumour-types from the International Cancer Genome Consortium/The Cancer Genome Atlas Pan-cancer Analysis of Whole Genomes (PCAWG)^43^ projects. We quantified ancestry associations in driver mutations, subclonal architecture, mutation timing and mutational signatures in almost all tumour-types, many linked to clinical phenotypes.

## Results

### Ancestry associations in Mutation Density and Timing

We separately analyzed TCGA and PCAWG data. These studies differ in molecular profiling technology, available annotation data, and populations sampled. We took the union of all TCGA tumours for pan-TCGA analyses, and the union of all PCAWG tumours for pan-PCAWG analyses. In addition to these pan-cancer analyses, we also examined each tumour-type for tumour-type-specific associations (**Table 1**). We used TCGA ancestry as previously imputed by Yuan *et al*^40^ and PCAWG ancestry data as previously reported^43^. We adapted a statistical approach previously applied to quantify sex-associations in cancer genomes^44^. Briefly, we first used univariate methods to identify putative associations within each non-European ancestry group: East Asian (EAN), African (AFR), American Indian and Alaska Natives (Admixed American; AMR), South Asian (SAN), and Other Ancestry (OA). European ancestry was used as the reference group because of its larger sample-size in these cohorts, which maximizes statistical power. An ancestry group was only studied in a tumour-type if its group sample size in that tumour-type was at least five (**Supplementary Table 1**). Putatively associated genomic features were then further modeled using multivariate regression to adjust for confounding clinico-epidemiologic factors such as sex, age, stage and tumour-type, and assessed for significance at a false discovery rate (FDR) threshold of 10% (**Methods**). Pan-cancer and tumour-type-specific models and variable specifications are presented in **Supplementary Table 1**.

We first asked whether genome-wide phenomena were associated with ancestry. We started with measure of genome instability, and initially focused on the burden of copy number alterations (CNAs), approximated by the proportion of the genome with a CNA (PGA)^45^. Using Mann-Whitney U-tests followed by linear regression (LNR), we identified significant associations between EAN ancestry and PGA in TCGA-hepatocellular cancer (LIHC: Δloc = 5.8%, 95%CI = 2.6 - 8.9%, adjusted LNR p = 6.0×10^−3^), TCGA-stomach and esophageal cancer (STES: Δloc = 5.7%, 95%CI = 1.6 - 9.9%, adjusted LNR p = 0.012), and pan-TCGA analyses (Δloc = 1.6%, 95%CI = 0.05 - 3.2%, adjusted LNR p = 0.012); PGA was associated with AFR ancestry in TCGA-head and neck cancer (HNSC: Δloc = 6.6%, 95%CI = 1.8 - 11% adjusted LNR p = 0.030), TCGA-endometrial cancer (UCEC: Δloc = 3.2%, 95%CI = 0.020 - 7.0% adjusted LNR p = 7.9×10^−3^) and pan-PCAWG analyses (Δloc = 8.8%, 95%CI = 4.3 - 13%, adjusted LNR p = 3.0×10^−3^; **Figure 1A**, **Supplementary Figure 1**). For all significant associations, PGA was higher in EAN- or AFR-derived tumours compared with tumours arising in individuals of EUR ancestry (**Figure 1B**, **Supplementary Figure 1**), indicating higher genome instability in the EAN- and AFR-derived tumours of these tumour-types.

**Figure 1.**
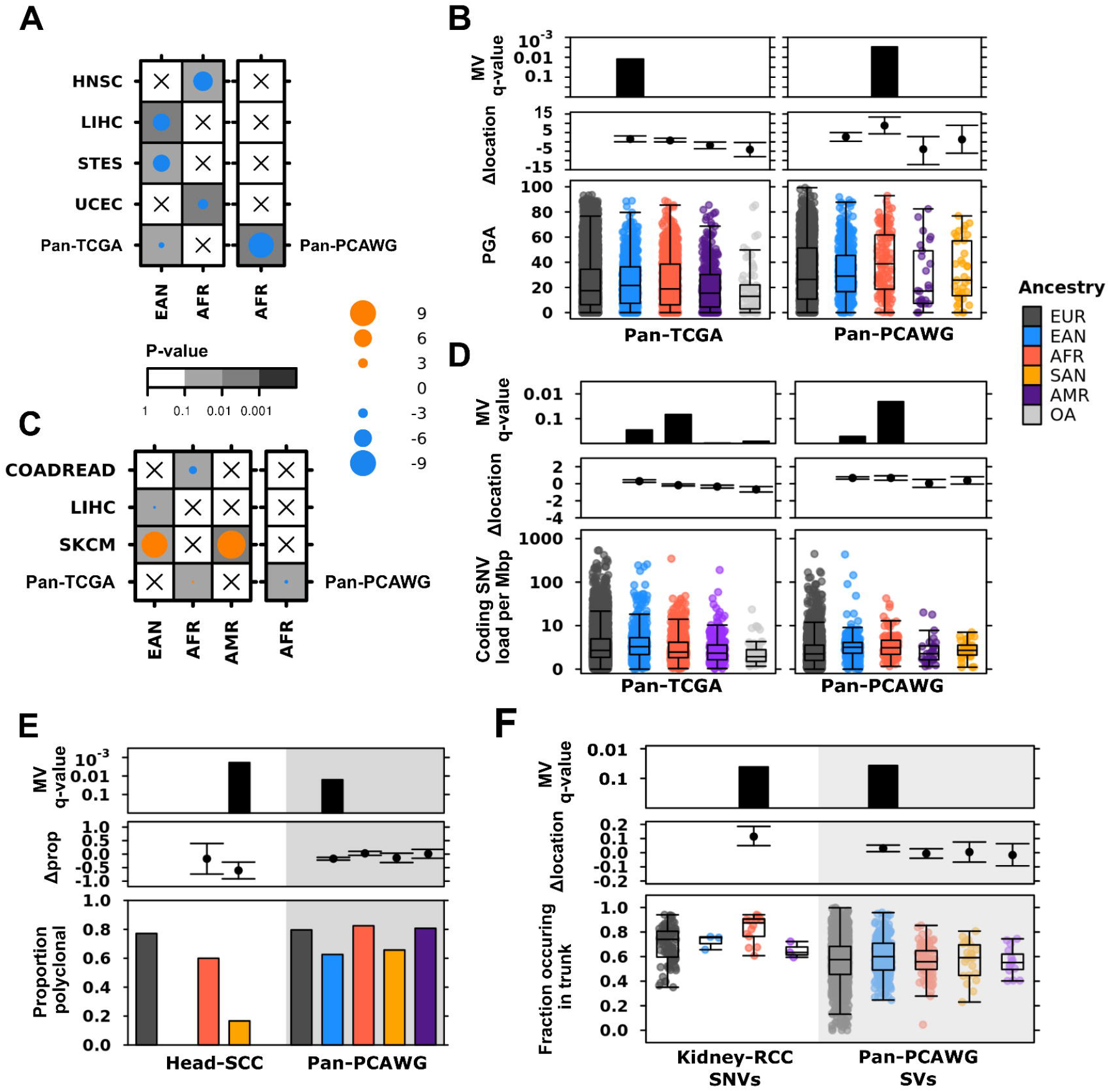
Ancestry associations in mutation density and evolutionary architecture. **(A)** Summary of associations between ancestry and percent genome altered (PGA) in TCGA and PCAWG tumours. The dot size and colour show the difference in location effect size estimate, and background shading indicate multiple-testing adjusted multivariate p-value. Only tumour-types with significant associations are shown. **(B)** Pan-TCGA and pan-PCAWG associations between ancestry and PGA, with the top row of barplots showing adjusted multivariate p-values, the middle row showing differences in location (mean and 95% confidence interval), and the bottom row of boxplots showing PGA per tumour. **(C)** Associations of ancestry with SNV density (SNVs/Mbp sequenced) in TCGA and PCAWG. Dot size and colour, and background shading have the same meaning and scale as Figure 1A. **(D)** pan-TCGA and pan-PCAWG associations between ancestry and SNV density, with adjusted multivariate p-values, differences in location, and PGA per tumour shown as in Figure 1B. **(E)** Differences in proportion of polyclonal tumours between ancestries with the top row showing adjusted multivariate p-value, middle row giving difference in proportion (mean and 95% confidence interval) and bottom row showing proportion of polyclonal tumours by ancestry. **(F)** Proportion of tumours with SNVs occurring in the truncal clone compared by ancestry in PCAWG kidney renal clear cell cancer, and truncal SVs in pan-PCAWG samples, with the same structure of rows as in Figure 1C. Tukey boxplots are shown with the box indicating quartiles and the whiskers drawn at the lowest and highest points within 1.5 interquartile range of the lower and upper quartiles, respectively.

Single nucleotide variation (SNV) density is an analogous measure to PGA, quantifying the burden of somatic SNVs in each Mbp of DNA sequenced. Again, we applied Mann-Whitney U-tests and linear regression to identify tumour-types where SNV density was associated with ancestry. We first assessed coding SNV density, as TCGA SNV data is based on whole exome sequencing (**Figure 1C**, **Supplementary Figure 1**). In TCGA-melanoma, both EAN (SKCM: Δloc = −8.3 SNVs/Mbp, 95%CI = −18 - −3.1 SNVs/Mbp, adjusted LNR p = 0.010) and AMR (Δloc = −9.0 SNVs/Mbp, 95%CI = −26 - −2.2 SNVs/Mbp, adjusted LNR p = 1.8×10^−3^) ancestries were associated with lower SNV density than the EUR reference. EAN-derived LIHC tumours (Δloc = 3.7 SNVs/Mbp, 95%CI = 0.1 – 0.67 SNVs/Mbp, adjusted LNR p = 1.8×10^−3^) and AFR-derived TCGA-colorectal (COADREAD: Δloc = 2.1 SNVs/Mbp, 95%CI = 0.53 - 4.1, adjusted LNR p = 0.068) tumours had higher SNV density than the EUR references. pan-TCGA, SNV density was lower in AFR-derived tumours (Δloc = −0.17 SNVs/Mbp, 95%CI = −0.30 - −0.067, adjusted LNR p = 0.068; **Figure 1D**). In contrast, coding SNV density was higher in AFR-derived pan-PCAWG tumours (Δloc = 0.68 SNVs/Mbp, 95%CI = 0.43-0.95, adjusted LNR p = 0.021). This difference in AFR-associated coding SNV density between pan-TCGA and pan-PCAWG data may highlight differences in included tumour-types and geographic differences in the populations sampled. Finally, we extended beyond the coding regions to examine non-coding and overall SNV density in the PCAWG whole genome sequencing data. These results closely matched those for coding SNV density; pan-PCAWG AFR-derived tumours had consistently lower SNV density regardless of the coding context (**Supplementary Table 2**).

We next focused on clonal architecture and mutation timing using data describing the evolutionary history of PCAWG tumours^46^. We tested whether monoclonal status might be ancestry-associated by comparing the proportions of tumours that were monoclonal, where all tumour cells are homogenous, clonal copies of one ancestral cell *vs* tumours that were polyclonal, which have multiple somatically distinct cells derived of different ancestral lineages. We used proportion tests followed by logistic regression (LGR) to identify ancestry-associations in monoclonal status. SAN-derived PCAWG-head and neck tumours were more frequently monoclonal than EUR-derived tumours (Head-SCC: Δproportion polyclonal = 0.60, 95%CI = 0.30-0.91, adjusted LGR p = 1.9×10^−3^; **Figure 1E**). In pan-PCAWG analysis, EAN-derived tumours were also more frequently monoclonal than EUR-derived tumours (Δproportion polyclonal tumours = 0.17, 95%CI = 0.12-0.22, adjusted LGR p = 0.016). Monoclonal tumours have previously been associated with better survival in several tumours types^47–49^, and a higher frequency of monoclonal tumours in SAN and EAN tumours might underlie some of the improved survival experienced by these ancestry groups in these tumour types.

Focusing only on polyclonal tumours, we investigated whether the time at which mutations accumulate during a tumour’s evolution might be associated with the ancestry of the patient it arises in. We compared how frequently SNVs, indels and structural variants (SVs) occurred as clonal mutations in the trunk or as subclonal ones in branches. In PCAWG-kidney renal clear cell cancer, tumours arising in AFR individuals had a higher proportion of clonal SNVs relative to those in EUR individuals (Kidney-RCC: Δloc = 0.11, 95%CI = 0.050 – 0.19, LGR p = 0.041; **Figure 1F**). Kidney cancer tumours arising in AFR individuals also had a higher proportion of truncal indels (Δloc = 0.11, 95%CI = 0.036 – 0.18, LGR p = 0.035; **Supplementary Table 2**). In pan-PCAWG tumours, EAN-derived tumours had a lower proportion of truncal SVs (Δloc = −0.03, 95%CI = −0.054 – −0.074, LGR p = 0.037; **Figure 1F**). Ancestry associations in monoclonal status and mutation timing suggest potential differences in the evolutionary histories of these SAN-, AFR- and EAN-derived tumours. Future investigations of tumour evolution using larger cohorts and multi-region sequencing are needed to validate and quantify these ancestry-associations.

### Ancestry Associations in Mutational Signatures

Differences in mutation density and timing suggest that the mutagenic processes affecting a tumour might be correlated with the ancestry of the patient, presumably primarily through differential environmental exposures associated with race and ethnicity. Mutational signatures based on the flanking sequence context of mutations can deconvolve characteristic mutational patterns that arise from specific mutagenic processes. We analysed three types of mutational signatures generated by the PCAWG project: 49 single base substitution (SBS), 11 doublet base substitution (DBS) and 17 small insertion and deletion (ID) signatures^50^. We also investigated SBS signatures for TCGA tumours. Each signature is thought to reflect a specific mutagenic process, though many are still of unknown aetiology^50,51^. For each signature, we examined both the proportion of signature-positive tumours as well as relative signature activity, quantified as the proportion of mutations attributed to each signature.

Ancestry-associated mutational signatures were identified in pan-PCAWG and pan-TCGA analyses, as well as in in four PCAWG and thirteen TCGA tumour-types (**Supplementary Table 2**). AFR-associations in mutational signatures occurred across several tumour-types (**Figures 2A, 2B**). In TCGA-lung adenocarcinoma, AFR-derived tumours had higher detection rates of SBS4, attributed to tobacco exposure (LUAD: Δproportion = 0.39, 95%CI = 0.23-0.55, adjusted LNR p = 5.5×10^−3^; **Figure 2A, Supplementary Figure 2**). Higher rates of SBS4 detection in AFR-derived lung adenocarcinoma despite reportedly comparable smoking rates between Black and White populations^52^ may reflect differences in nicotine metabolism^53^, elevated use of mentholated cigarettes in Black populations^52,54^ or simple selection bias in the cohort of lung adenocarcinomas studied in TCGA. In PCAWG-breast cancer, SBS3 occurred more frequently in AFR-derived samples (Breast-AdenoCA: Δproportion = 0.33, 95%CI = 0.13-0.54, adjusted LNR p = 0.097). SBS3 is attributed to defective homologous recombination (HR) repair of double stranded breaks. Higher SBS3 occurrence in AFR breast tumours may be due to more frequent triple-negative breast cancer, which exhibit high rates of defective HR repair^55^. Finally, AFR-derived TCGA-endometrial cancers had higher detection rates of SBS2 (UCEC: Δproportion = 0.26, 95%CI = −0.0081-0.54, adjusted LNR p = 5.2×10^−2^). and SBS13 (Δproportion = 0.31, 95%CI = 0.030-0.59, adjusted LNR p = 0.011), which are both attributed to the activity of AID/APOBEC cytidine deaminases and have been previously associated with progression from primary to metastatic disease^56^. AFR-associations in relative signature activity were detected in two signatures of unknown aetiology (**Figure 2B**).

**Figure 2.**
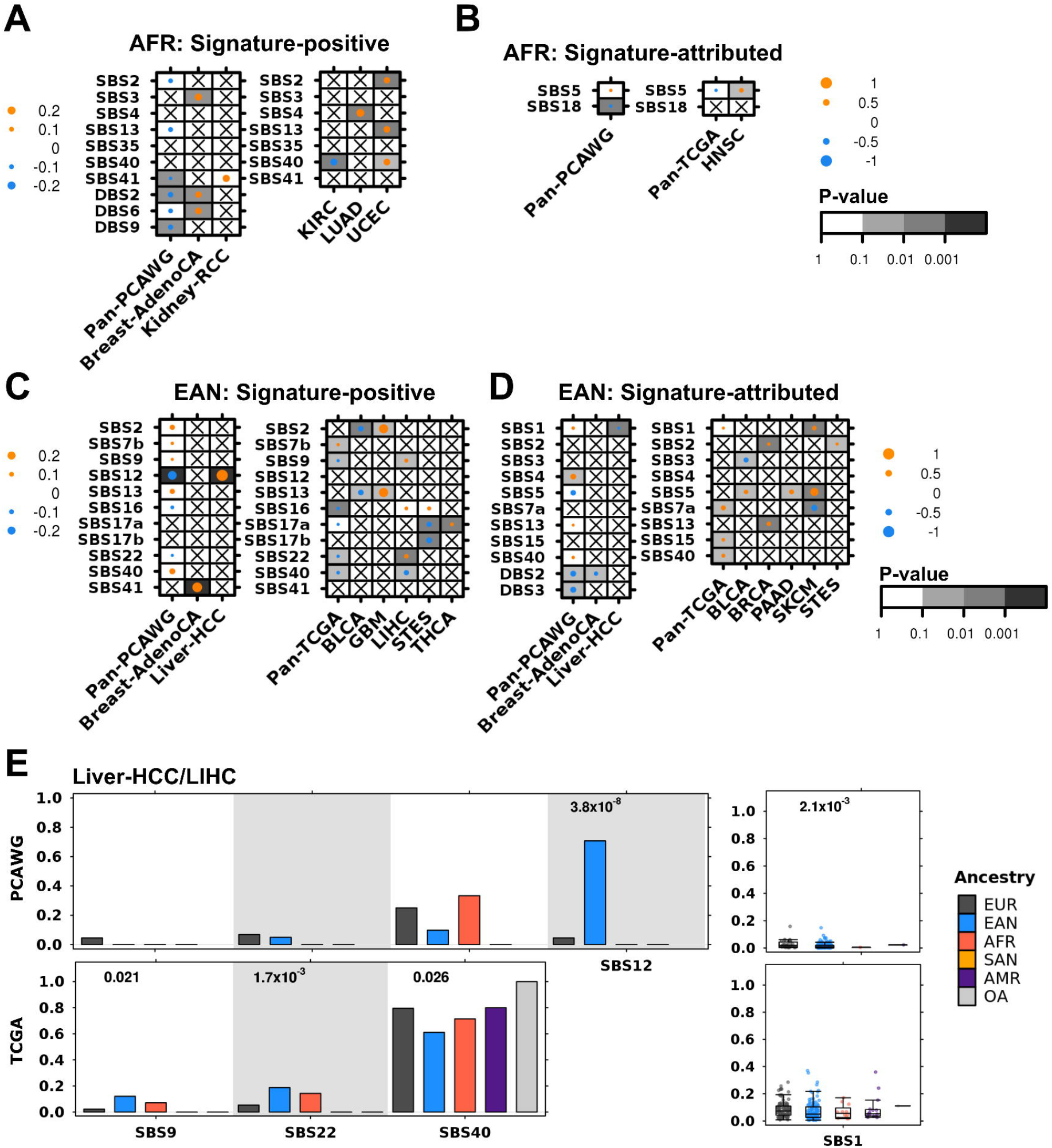
Ancestry-associations in mutational signatures. **(A)** Summary of associations between AFR ancestry and the proportion of signature-positive tumours. Here dot size and colour indicate the differences in proportion between AFR and EUR tumours, and the background shading gives multiple-testing adjusted multivariate p-values. PCAWG data is on left and TCGA on right. **(B)** Similarly, the summary of associations between AFR ancestry and relative signature activity, with dot size showing difference in location estimates and background indicating multiple-testing adjusted linear regression p-values. **(C)** Summary of associations between EAN ancestry and signature positive tumours, as for Figure 2A. **(D)** Summary of associations between EAN ancestry and relative signature activity, as in **(B)**. **(E)** Ancestry-associated differences in hepatocellular cancer, compared between PCAWG and TCGA data. Barplots show frequency of signature detection in each ancestry group. Tukey boxplots (as described in Figure 1) show relative signature activity as proportion of mutations attributed to each signature.

Of the ancestries analysed, the largest number of significant associations were detected with EAN ancestry (**Figures 2C, 2D**). In hepatocellular cancer, signatures were EAN-associated in both proportion positive and relative activity across both PCAWG (Liver-HCC; **Figure 2E, top**) and TCGA (**Figure 2E, bottom**) data. EAN-derived Liver-HCC tumours had higher SBS12 detection frequency (Δproportion = 0.66, 95%CI = 0.57-0.76, adjusted LNR p = 3.8 × 10^−8^) and lower relative SBS1 activity (Δloc = 0.0067, 95%CI = 0.0019-0.015, adjusted LNR p = 2.1×10^−3^) compared with EUR hepatocellular tumours; SBS12 was not described in TCGA-hepatocellular cancer data, and decreased SBS1 activity in EAN-derived TCGA samples was not statistically significant after multivariable adjustment. In TCGA-hepatocellular tumours, EAN-derived tumours showed higher rates of SGS9 signature detection (Δproportion = 0.098, 95%CI = 0.033-0.16, adjusted LNR p = 0.021), SBS22 (Δproportion = 0.13, 95%CI = 0.054-0.22, adjusted LNR p = 1.7×10^−3^) and SBS40 (Δproportion = −0.18, 95%CI = -−0.30 - 0.073, adjusted LNR p = 0.026). Intriguingly, the TCGA EAN-associations in SBS9 and SBS22 were not reflected in PCAWG data despite sufficient group sample sizes (**Figure 2E; Supplementary Table 1**). SBS9 is attributed to mutations induced during replication by DNA polymerase η and SBS22 to aristolochic acid exposure. These contrasting results between PCAWG and TCGA data may be due to ethnic and geographic differences between the datasets: PCAWG hepatocellular tumours were primarily from Japanese and French patients, while TCGA tumours are from US patients.

Other ancestry-associated mutational signatures include higher relative activity of ID2 in SAN-derived PCAWG-head and neck tumours (**Supplementary Figure 2, Supplementary Table 2**). ID2 is attributed to slippage of the template strand during DNA replication and is thought to be associated with DNA mismatch repair deficiency. SBS5 was detected at lower rates in AMR TCGA-bladder cancer (BLCA: Δproportion = −0.15, 95%CI = −0.41 - 0.10, adjusted LNR p = 0.012; **Supplementary Figure 2, Supplementary Table 2**) and SBS16 at higher rates in AMR-derived TCGA-lower grade glioma (LGG: Δproportion = 0.079, 95%CI = −0.054 - 0.21, adjusted LNR p = 0.029; **Supplementary Figure 2**). AMR-derived LGG tumours also had higher relative activity of AID/APOBEC-attributed SBS13 (Δloc = 0.0068, 95%CI = 0.030-0.012, adjusted LNR p = 6.0×10^−3^, **Supplementary Figure 2**), suggesting a greater role of these enzymes in lower grade gliomas of individuals of AMR ancestry. Thus ancestry-associated mutational signatures were detected across a range of endogenous and exogenous mutational processes. Most ancestry-associated signatures are of unknown aetiology and elucidation of the biological processes underlying these signatures may help determine if these are true ancestry-associations or confounding from other environmental differences in the cohorts.

### Ancestry-associations in CNA Differences are Associated with Transcriptomic Changes

After identifying ancestry-associated differences in genome-wide phenomena and mutational signatures, we focused on chromosome segment and gene-level events. We compared the proportions of copy number losses and copy number gains for each ancestry group compared with the EUR reference group using proportion tests, and then adjusted for confounding factors using multivariate logistic regression. As with prior analyses, we analysed TCGA and PCAWG concurrently and separately. Ancestry-associated CNAs were identified in 13 TCGA and 3 PCAWG cancer types, as well as in both pan-TCGA and pan-PCAWG analyses (**Figure 3A**). Pan-TCGA, 602 genes were in EAN-associated CNAs and 2,787 genes in AFR-associated CNAs. Some differences in CNA frequency were as large as 10% (**Figure 3B, Supplementary Table 3-4**). In pan-PCAWG analysis, 288 genes were in EAN-associated CNAs, 5,589 genes in AFR-associated CNAs, and 437 genes in SAN-associated CNAs, with frequency differences of up to 20% (**Supplementary Figure 3**, **Supplementary Tables 3-4**).

**Figure 3.**
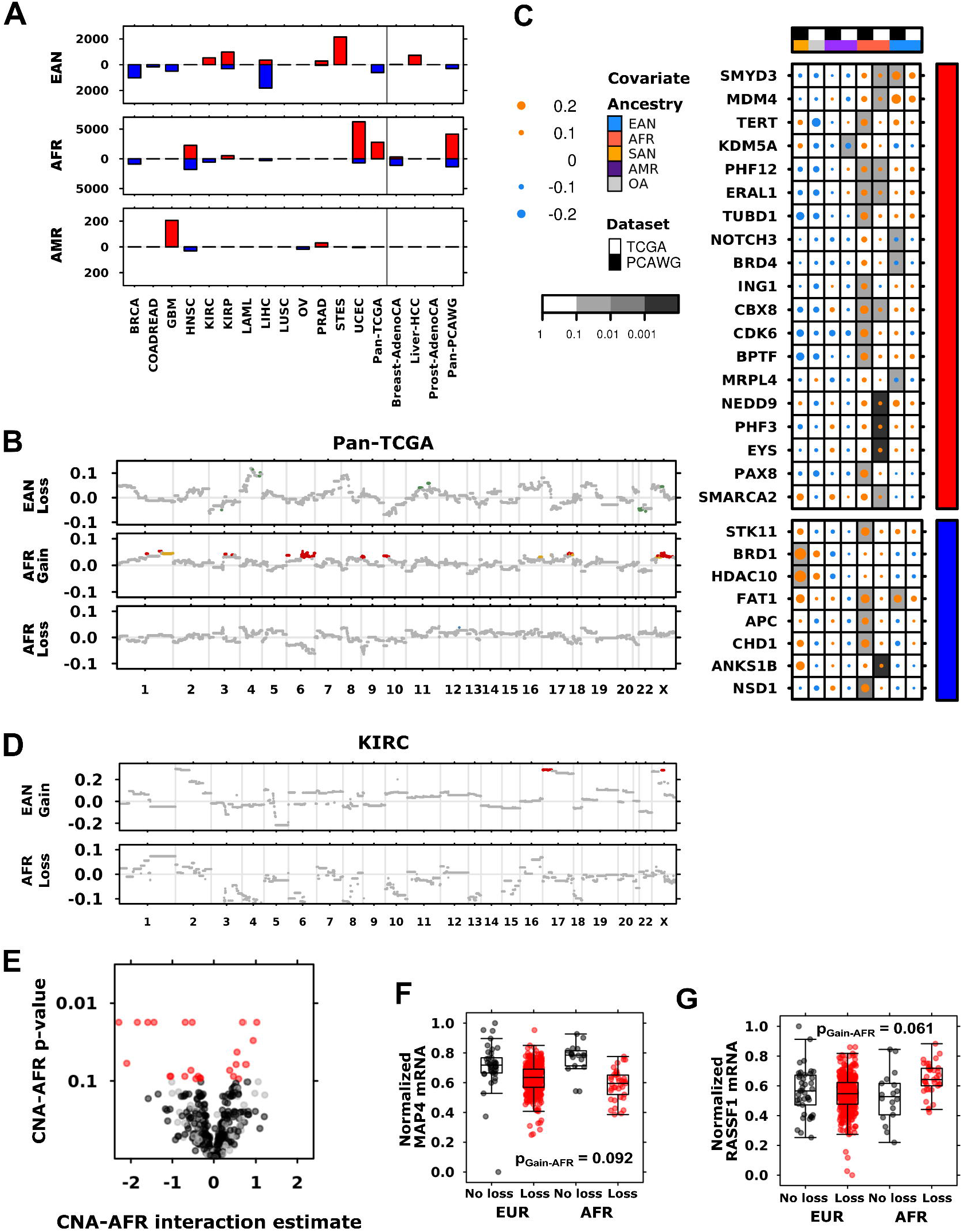
CNA-Ancestry associations are associated with altered RNA abundance. **(A)** Summary of all detected ancestry-associated CNAs with numbers of gains (above x-axis) and losses (below x-axis) identified in each tumour context. Only tumour-types with at least one significant event are shown. **(B)** Pan-cancer ancestry-associations in CNAs for TCGA data. Each plot shows the logistic regression coefficient estimate (for age analyses) or difference in proportion (for ancestry analyses) for the indicated variable and CNA type. Dot colour indicates statistical significance, where red (copy number gain) and blue (copy number loss) show adjusted p < 0.05 and yellow (gain) and green (loss) show whether the multiple-testing adjusted p < 0.1 threshold is met. **(C)** Summary of ancestry-associated pan-cancer CNA drivers. Both TCGA and PCAWG findings are shown, and dot size indicates the effect-size as a proportion difference. Background shading shows multiple-testing adjusted multivariate p-values. The covariate to the right shows copy number gain drivers in red and loss drivers in blue. **(D)** EAN- and AFR-associations in TCGA kidney clear cell cancer CNAs are associated with **(E)** changes in mRNA abundance. The adjusted p-value is plotted against the coefficient of the CNA-age interaction for mRNA abundance, with each point representing a gene. Black dots show significant associations between mRNA and CNA; red dots show significant CNA-ancestry interactions. **(F)***RASSF1* and **(G)** MAP4 mRNA abundance changes between copy number loss (red) or no loss (black) in tumours of EUR and AFR ancestry. Adjusted CNA-AFR interaction p-value is shown. Tukey boxplots are depicted, as described in Figure 1.

To determine whether cancer drivers were affected by ancestry-associated CNAs, we focused on a subset of 133 genes altered by driver CNAs^57^ (**Figure 3C**). There were eight pan-TCGA and 20 pan-PCAWG AFR-associated genes, including higher frequency of *CBX8* gain (PCAWG Δproportion = 0.16, 95%CI = 0.068 – 0.26, adjusted LGR p = 0.020; TCGA Δproportion = 0.043, 95%CI = 0.012 – 0.074, adjusted LGR p = 0.060) and *SMARCA* gain (PCAWG Δproportion = 0.12, 95%CI = 0.042 – 0.20, adjusted LGR p = 0.082; TCGA Δproportion = 0.030, 95%CI = 0.0058 – 0.055, adjusted LGR p = 0.013). One gene, *FAT1* was more frequently lost in tumours derived of EAN individuals (PCAWG Δproportion = 0.15, 95%CI = 0.096 – 0.20, adjusted LGR p = 0.080; TCGA Δproportion = 0.096, 95%CI = 0.055 – 0.14, adjusted LGR p = 0.14). Similarly, ancestry-associated driver CNAs were identified in 16 TCGA and 5 PCAWG cancer types (**Supplementary Tables 3-4**).

CNAs change the dosage of affected genes and can lead to transcriptome changes^58^. We sought to determine whether ancestry-associated CNAs have such downstream mRNA associations. Using TCGA mRNA abundance data, we analysed the mRNA of genes contained in ancestry-associated CNAs using models that incorporated the ancestry group of interest, copy number status, and the interaction between copy number status and ancestry. These models allowed us to identify mRNA abundance changes associated with the CNA itself, as well as changes where the effect of the copy number depended on the ancestry of the patient in which the tumour arose. We also adjusted for tumour purity as estimated by study pathologists in all mRNA analyses.

In TCGA-kidney clear cell cancer, 559 genes were present in AFR-associated losses (**Figure 3D**). Using the approach described above, we examined mRNA abundance for each of these genes as a function of AFR ancestry, copy number loss status, and their interaction. Of the 559 genes, copy number loss was significantly associated with changes in mRNA abundance for 316 genes (57%; **Figure 3E**, **black points**). A further 24 genes (4.3%) had significant interactions between copy number loss status and AFR ancestry (**Figure 3E**, **red points**). For some genes, this significant interaction was due to differences in the magnitude of mRNA abundance change: for example, loss of the microtubule-associated gene *MAP4* is associated with decreased *MAP4* mRNA abundance in both EUR and AFR tumours, but the decrease in mRNA abundance is greater in AFR than in EUR tumours (**Figure 3F**). For other genes, the significant interaction indicates a difference in direction: loss of the tumour suppressor *RASSF1* is associated with a slight decrease in *RASSF1* mRNA abundance for EUR-derived tumours, but an increase in abundance for AFR-derived tumours (**Figure 3G**). Thus copy number loss of *RASSF1* is not only less frequent in AFR-derived tumours (Δproportion = −0.24 95%CI = −0.38 - −0.10, adjusted LNR p = 1.3×10^−6^), its effect on mRNA abundance is contrary to what is usually observed in EUR-derived kidney tumours.

Repeating this mRNA analysis for all TCGA tumour-types with ancestry-associated CNAs, 5-53% of genes affected by EAN- or AFR-associated CNAs were significantly associated with changes in mRNA abundance (**Supplementary Table 5**). We also identified additional mRNA where the changes in abundance were dependent on the interaction between CNA and ancestry (**Supplementary Figure 3, Supplementary Table 5**). Significant CNA-EAN interactions were found in breast cancer (4 genes), kidney renal clear cell (5 genes) and papillary cell cancers (9 genes), liver cancer (8), lung squamous cell cancer (LUSC: 2), prostate cancer (PRAD: 13), and stomach and esophageal cancer (32 genes). There were also significant CNA-AFR interactions for UCEC (3; **Supplementary Figure 3**). Thus, CNA frequency is associated with ancestry, and ancestry-associated CNAs are associated with changes in the transcriptome.

### Ancestry-associations in Gene-Level SNVs

Finally, we asked whether specific genes might be mutated by SNVs at different frequencies between different ancestry groups. In TCGA data, we applied a recurrence filter and removed genes that had SNVs in <1% of tumours for each tumour-type. In PCAWG data, we focused on a set of drivers^59^ that includes both coding and non-coding elements, as well as SNVs in mitochondrial DNA (mtDNA). Ancestry-associations were identified in 15 TCGA tumour-types and four PCAWG tumour-types (**Figure 4A**). Across pan-TCGA tumours, there were nine genes that were mutated with SNVs more frequently in EAN-derived samples including *FGFR3 (*Δproportion = 0.022, 95%CI = 0.054-0.039, adjusted LGR p = 0.016; **Supplementary Table 6**). In pan-PCAWG tumours, SNVs in the coding regions of the tumour suppressors *FBXW7* (Δproportion = 0.066, 95%CI = 0.014-0.12, adjusted LGR p = 0.030) and *TP53* (Δproportion = 0.18, 95%CI = 0.087-0.28, adjusted LGR p = 0.044) occurred more frequently in tumours arising in AFR individuals (**Figure 4B, left**). *TERT* promoter SNVs were more frequent in tumours arising in SAN individuals (Δproportion = 0.17, 95%CI = 0.012-0.33, adjusted LGR p = 0.024).

**Figure 4.**
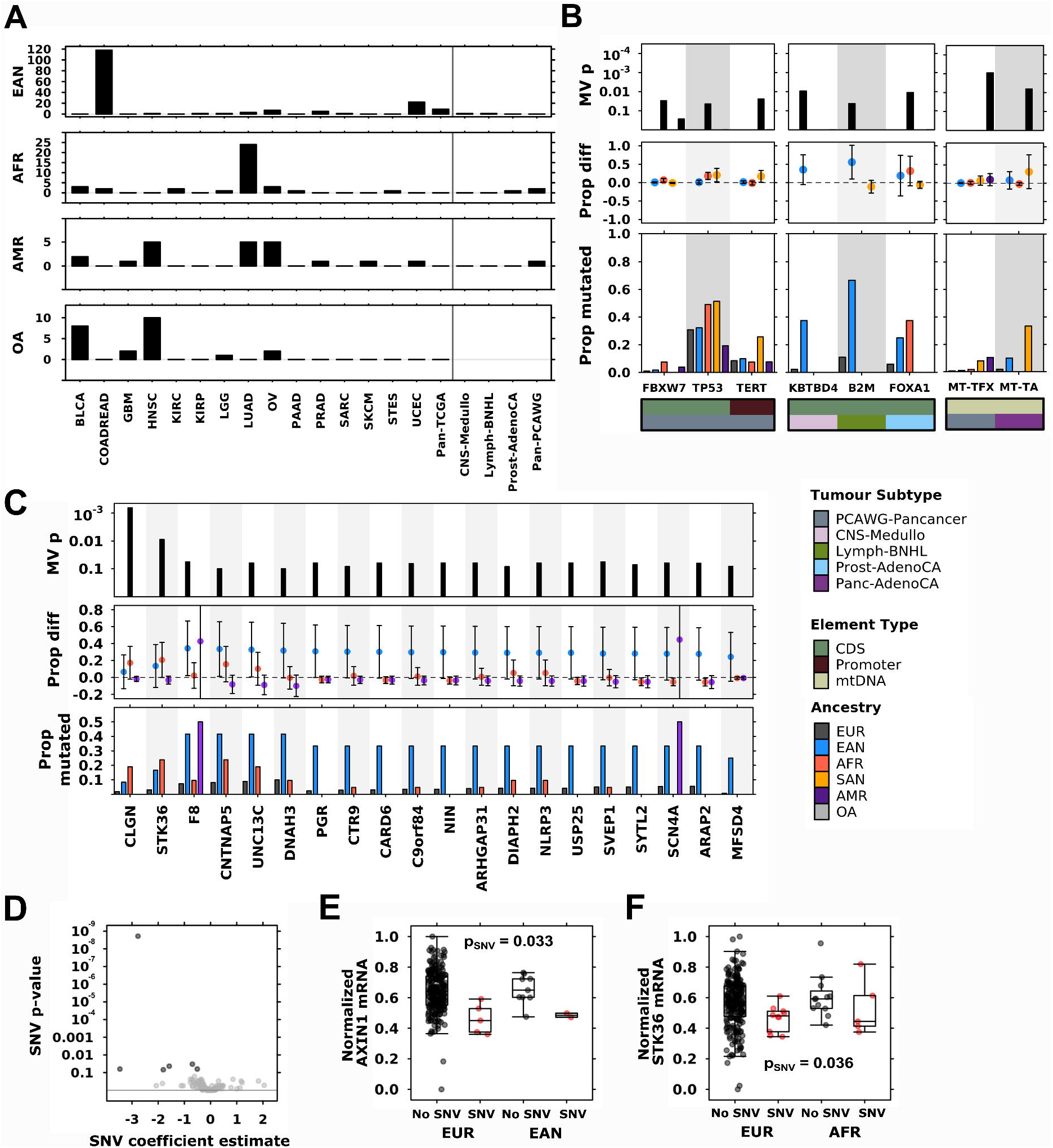
Ancestry-associations in gene-level SNV mutation frequency. **(A)** Summary of all detected ancestry-associated SNVs found in each cancer type. Only tumour-types with at least one significant event shown. **(B)** pan-PCAWG and PCAWG tumour-type-specific ancestry-associations in driver and mitochondrial SNV frequency with the top showing adjusted multivariate p-values, middle showing difference in proportion, and bottom showing proportion of tumours with mutated gene per ancestry group. Covariate bars indicate tumour and element type context of each mutation, where coding sequence is abbreviated to CDS and mitochondrial DNA is mtDNA. **(C)** The 20 SNVs most associated with AFR and EAN ancestry in the TCGA colorectal and renal cancer (COADREAD) dataset. **(D)** Differential mRNA abundance associated with EAN-associated SNVs in TCGA COADREAD. Each point represents a gene with black dots showing significant associations between mRNA and SNV. **(F)** *AXIN1* and **(G)** STK36 mRNA abundance changes between copy number loss (red) or no loss (black) compared by ancestry. Adjusted SNV term p-value is shown. Tukey boxplots are depicted, as described in Figure 1.

We also identified EAN-associated SNVs in PCAWG medulloblastoma and PCAWG non-Hodgkin’s lymphoma (**Figure 4B, middle**). SNVs in both *KBTBD4* (CNS-Medullo: Δproportion = 0.35,95%CI = −0.050 −0.75, adjusted LGR p = 9.9×10^−3^) and *B2M* (Lymph-BNHL: Δproportion = 0.56,95%CI = 0.086-1, adjusted LGR p = 0.043) occurred more frequently in EAN-derived tumours than EUR. In PCAWG-prostate cancer, SNVs in *FOXA1* were more frequently in tumours derived of AFR individuals (Prost-AdenoCA: Δproportion = 0.31,95%CI = −0.088-0.72, adjusted LGR p = 0.012). The frequency of mitochondrial SNVs were also associated with ancestry (**Figure 4B, right**): pan-PCAWG, SNVs in *MT-TFX* occurred more frequently in AMR-derived tumours (Δproportion = 0.098,95%CI = −0.066-0.26, adjusted LGR p = 9.0×10^−4^), and *MT-TA* SNVs were more frequent in SAN-derived pancreas tumours (Δproportion = 0.31,95%CI = −0.15-0.78, adjusted LGR p = 6.5×10^−5^). *MT-TFX* is a mitochondrially encoded transcription factor binding site, and *MT-TA* encodes a transfer RNA for alanine. Mutations in mtDNA could have far-reaching downstream effects. For example, mutations in *MT-TA* could result in less efficient protein synthesis, leading to differences in the tumour proteome.

Across all TCGA tumour-types, we identified 159 EAN-associations, 37 AFR-associations, and 23 AMR-associations. These included genes in KIRC such as *UBR5* which was mutated by SNVs more frequently in AFR-derived tumours (Δproportion = 0.14,95%CI = −0.038-0.33, adjusted LGR p = 9.8×10^−3^), and in COADREAD, where *CARD6* (COADREAD: Δproportion = 0.30,95%CI = −0.0075-0.61, adjusted LGR p = 0.062; **Figure 4C**) and *CKAP2* (Δproportion = 0.24,95%CI = −0.049-0.53, adjusted LGR p = 0.062) SNVs were more frequent in EAN-derived TCGA-colorectal tumours (**Supplementary Table 6**). The majority of ancestry-associated SNVs were identified in COADREAD, which had 118 EAN-associations and two AFR-associations in COADREAD gene-level SNVs (**Figure 4C**).

Similar to our CNA analyses, we next investigated mRNA abundance to determine whether ancestry-associated SNVs might also be associated with changes in the transcriptome. We used the same approach as previously applied in our CNA-transcriptome analyses, using a model that included SNV status, ancestry, and the interaction between SNV and ancestry. Despite low statistical power due to small group sizes, several ancestry-associated SNVs were associated with changes in mRNA abundance. In COADREAD, six genes that were more frequently mutated with SNVs in EAN-derived tumours were also associated with decreased mRNA abundance (**Figure 4D, Supplementary Table 6**), including *AXIN1* (**Figure 4E**). Similarly SNVs in *STK36,* which occurred more frequently in AFR-derived COADREAD (Δproportion = 0.21,95%CI = −0.00058-0.41, adjusted LGR p = 8.8×10^−3^), was also associated with decreased mRNA abundance in tumours derived of both EUR and AFR individuals (**Figure 4F**). We identified differential mRNA abundance associated with ancestry-associated SNVs in TCGA-bladder cancer (BLCA) and UCEC (**Supplementary Table 6**). Unlike in our CNA analysis, we did not find mRNA changes that were dependent on the interaction between SNV and ancestry.

## Discussion

Our analysis of TCGA and PCAWG data revealed ancestry-associations across all genomic features studied, from genome-wide phenomena to gene-level events. These associations occur at both the pan-cancer level and in specific tumour-types. Ancestry is associated with mutation density, measures of tumour evolution, and with mutational signatures associated with oncogenic processes. Gene-level CNAs and SNVs not only occurred at different frequencies in different ancestries, they were also associated with differential mRNA abundance: in some cases, the effect of a CNA on mRNA abundance was dependent on the ancestry of the patient in which the tumour arose. Together, these results suggest that ancestry influences the progression of a tumour and that ancestry-associated genomic events have potential functional significance.

Differences between the TCGA and PCAWG datasets allowed us to investigate them in parallel and orthogonal ways. We leveraged the deeper clinical annotation and larger samples sizes of TCGA whole exome sequencing and array-based data to adjust for more confounding variables using more complex models. In contrast, the whole genome sequencing data from PCAWG allowed us to investigate a broader range of genomic features such as clonal architecture and non-coding drivers. TCGA and PCAWG data also represent different geographic populations: while TCGA tumours were largely derived of North American patients, PCAWG tumours were from patients at multiple international sites. As a result, the distributions of ancestry groups differ between TCGA and PCAWG data (**Table 1**), and many PCAWG tumour-types were excluded from tumour-type-specific analysis due to insufficient sample size. Poor agreement between TCGA and PCAWG results are therefore related to three major factors: vastly different sample sizes and ancestry group sizes affecting statistical power; geographical differences in sampling; and differences in molecular profiling technologies.

As with other ancestry- and race-associated differences in cancer burden, the causes of ancestry-associated genomic events are multifactorial and interacting. For example, our analysis of mutational signatures revealed the differing impacts of both endogenous and exogenous mutagens on tumours of different ancestries. One such difference was in the increased detection rates of defective homologous recombination repair in breast tumours derived of individuals of African ancestry. However, whether this increase is due to differences in inherited predisposition^60^, hormonal differences^61^, environmental exposure, or a combination of these and other variables is uncertain. Ancestry-associations in cancer genomes likely arise from a combination of biological, lifestyle, and environmental factors. We used imputed ancestry, which best approximates the line of descent for an individuals’ genetic material. However, without accounting for factors highly correlated with ancestry, such as race, socioeconomic status and quality of healthcare^62,63^, we cannot fully disentangle contributing factors and definitively attribute these differences to biology.

Ancestry undoubtedly influences the molecular presentation of a tumour. Despite low sample sizes, we have identified differences in the density, frequency, and transcriptional consequence of both copy number and nucleotide changes. The results we present are likely an underestimation of the full landscape of ancestry-associated somatic changes in cancer, and this is due in large part to poor representation of non-European ancestries in TCGA, PCAWG, and other cancer profiling studies^64^. To fully describe ancestry-associations, future genomic studies must include diverse representation across multiple ancestry groups and include deep and complete annotation to facilitate the control of confounding variables. Through identifying differences in the cancer genomes between individuals of different ancestries, we can better understand how they arise and leverage them to improve personalized therapy strategies.

## Online Methods

### Data acquisition & Processing

Genome-wide somatic copy-number, somatic mutation, and mRNA abundance profiles for the Cancer Genome Atlas (TCGA) datasets were downloaded from Broad GDAC Firehose (https://gdac.broadinstitute.org/), release 2016-01-28. For mRNA abundance, Illumina HiSeq rnaseqv2 level 3 RSEM normalised profiles were used. Genes with >75% of tumours having zero reads were removed from the respective dataset. GISTIC v2 (13) level 4 data was used for somatic copy-number analysis. mRNA abundance data were converted to log_2_ scale for subsequent analyses. Mutational profiles were based on TCGA-reported MutSig v2.0 calls. All pre-processing was performed in R statistical environment (v3.1.3). Genetic ancestry imputed by Yuan *et al*. was downloaded from The Cancer Genetic Ancestry Atlas (http://52.25.87.215/TCGAA).

PCAWG WGS data were downloaded from the PCAWG consortium with pre-processing as performed by the consortium^43^. Individual datasets are available at Synapse (https://www.synapse.org/), denoted with synXXXXX accession numbers (*i.e.* Synapse IDs). These datasets are mirrored at https://dcc.icgc.org. Tumour histological classifications were reviewed and assigned by the PCAWG Pathology and Clinical Correlates Working Group (annotation version 9; syn10389158, syn10389164). Ancestry imputation was performed using an ADMIXTURE-like algorithm based on germline SNP profiles determined by whole-genome sequencing of reference sample (syn4877977). The consensus somatic SNV and indel (syn7357330) file covers 2778 whitelisted samples from 2583 donors. Driver events were called by the PCAWG Drivers and Functional Interpretation Group (syn11639581). Consensus CNA calls from the PCAWG Structural Variation Working Group were downloaded in VCF format (syn8042988). Subclonal reconstruction was performed by the PCAWG Evolution and Heterogeneity Working Group (syn8532460). SigProfiler mutation signatures were determined by the PCAWG Mutation Signatures and Processes Working Group for single base substitution (syn11738669), doublet base substitution (syn11738667) and indel (syn11738668) signatures. Signatures data for TCGA, non-PCAWG WGS and non-TCGA WXS samples were downloaded from Synapse (syn11804040).

We used TCGA data on 10,212 distinct patients with 23 cancer types. PCAWG data was from 2,562 distinct patients with 29 cancer types. Cancer types with no age information or insufficient variability in ancestry annotation were excluded from analysis. TCGA genetic ancestry describes a five-category variable (European American, East Asian American, African American, Admixed American and Other Ancestry). PCAWG ancestry describes a five-category variable (European, East Asian, African, South Asian and Admixed American; **Supplementary Table 1**).

### General Statistical Framework

For each genomic feature of interest, we used univariate two-sided non-parametric tests followed by false discovery rate (FDR) adjustment to identify candidate ancestry-associations (q < 0.1). These were followed with multivariate modeling to account for potential confounders using tumour-type-specific models. We use EUR ancestry as the reference group for all analyses to maximize statistical power.

**Supplementary Table 1** gives the specific variables included for each cancer type. These were selected based on data availability (<15% missing), variability (at least two levels) and collinearity (as assessed by variance inflation factor). Discrete data was modeled using logistic regression (LGR). Continuous data was first transformed using the Box-Cox family and modeled using linear regression (LNR). The Box-Cox family of transformations is a formalized method to select a power transformation to better approximate a normal-like distribution and stabilize variance. We used the Yeo-Johnson extension to the Box-Cox transformation that allows for zeros and negative values^65^. FDR adjustment was performed for p-values for ancestry variable significance estimates, and a threshold of 10% used to select candidates. A summary of all results is presented in **Supplementary Table 1**. We present 95% confidence intervals for all tests.

### Mutation Density

Performed for both TCGA and PCAWG data. Overall SNP mutational density per calculated per patient was calculated as the count of SNVs scaled to SNVs/Mbp. Coding mutation prevalence only considers the coding regions of the genome, while noncoding prevalence considers only noncoding regions. TCGA mutation density is coding mutation prevalence. Mutation density was compared between ancestries using two-sided Mann-Whitney U-tests using European ancestry as the reference group (*e.g.* EAN *vs.* EUR, AFR *vs.* EUR, *etc*.). Comparisons with univariate q-values meeting an FDR threshold of 10% were analyzed using linear regression to adjust for tumour subtype-specific variables. Mutation density analysis was performed separately for each mutation context, with pan-cancer and tumour subtype p-values adjusted together. Full mutation density results are in **Supplementary Table 2**.

### Genome instability

Performed for both TCGA and PCAWG data. Genome instability was calculated as the percentage of the genome affected by copy number alterations. The number of base pairs for each CNA segment was summed to obtain the total number of base pairs gained and lost in at least one allele. This total was scaled by the number of bases in the human genome reference to obtain the proportion of the genome with a CNA (PGA). Genome instability was compared between ancestries using two-sided Mann-Whitney U-tests for both pan-cancer and tumour-type specific analysis. Comparisons with univariate q-values meeting an FDR threshold of 10% were then subject to linear regression to adjust for tumour subtype-specific variables. Genome instability analysis was performed separately for each mutation context, with pan-cancer and tumour subtype p-values adjusted together. Full mutation density results are in **Supplementary Table 2**.

### Clonal structure and mutation timing analysis

Performed for PCAWG data only. Subclonal structure data was discretized into monoclonal (one cluster) *vs.* polyclonal (more than one cluster). The proportion of polyclonal tumours was calculated for each ancestry. These proportions were compared with two-sided proportion tests and univariate FDR-adjusted p-values used to identify putatively sex-associated clonal structure. Candidates from this analysis were then subject to logistic regression to control for confounders, with a multivariate q-value threshold of 0.1 used to identify statistically significant ancestry-associations with clonal structure.

Mutation timing data classified SNVs, indels and SVs into clonal (truncal) or subclonal groups. The proportion of truncal variants was calculated for each mutation type (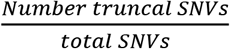, *etc.*) to obtain proportions of truncal SNVs, indels and SVs for each tumour. These proportions were compared between ancestries using two-sided Mann-Whitney *U*-tests. Univariate p-values were FDR adjusted to identify putatively ancestry-associated mutation timing. Linear regression was used to adjust for confounding factors and a multivariate q-value threshold of 0.1 was used to determine statistically significant ancestry-associated mutation timing. The mutation timing analysis was performed separately for SNVs, indels and SVs. All results for clonal structure and mutation timing analyses are in **Supplementary Table 2**.

### Mutational Signatures analysis

Performed for both TCGA and PCAWG data. For each signature, we compared the proportion of tumours with any mutations attributed to the signatures (“signature-positive”) using two-sided proportion tests to identify univariately significant ancestry-associations. Signatures with putative ancestry-associations were further analysed using multivariable logistic regression. We also compared relative signature activity by performing Mann-Whitney U-tests to compare the proportions of mutations attributed to each signature. Following these two-sided tests, candidate sex-associated signatures were subject to multivariable linear regression after Box-cox adjustment, as outlined above. Signatures not detected in a tumour subtype were omitted from analysis for that tumour subtype. All results for clonal structure and mutation timing analyses are in **Supplementary Table 2**.

### Genome-spanning CNA analysis

Performed for both TCGA and PCAWG data. Adjacent genes whose copy number profiles across patients were highly correlated (Pearson’s *r* > 95%) were binned. The copy number call for each patient was taken to be the majority call across all genes in each bin. Copy number calls were collapsed to ternary (loss, neutral, gain) representation by combining loss groups (mono-allelic and bi-allelic) and gain groups (low and high). Two-sided proportion tests were used to identify univariate ancestry-associated CNAs. After identifying candidate pan-cancer univariately significant genes, multivariate logistic regression was used to adjust ternary CNA data for tumour-type-specific variables. The genome-spanning analysis was performed separately for losses and gains for each tumour subtype. All CNA results are in **Supplementary Tables 3-4**.

### Genome-spanning SNV analysis

Performed for TCGA data. We focused on genes mutated in at least 1% of patients. Mutation data was binarized to indicate presence or absence of SNV in each gene per patient. Proportions of mutated genes were compared between ancestry groups using two-sided proportions tests for univariate analysis. False discovery rate correction was used to adjust p-values with q < 0.1 as a threshold for multivariate logistic regression.

### Driver Event Analysis

Performed for PCAWG data. We focused on driver events described by the PCAWG consortium^59^. Driver mutation data was binarized to indicate presence or absence of the driver event in each patient. Proportions of mutated genes were compared between ancestries using two-sided proportions tests. A q-value threshold of 0.1 was used to select genes for further multivariate analysis using binary logistic regression. FDR correction was again applied and genes with significant pan-cancer ancestry terms were extracted from the models (q-value < 0.1). Driver event analysis was performed separately for pan-cancer analysis and for each tumour subtype. All SNV and driver event analysis results are in **Supplementary Table 6**.

### mRNA abundance analysis

Performed for TCGA data. Genes in CNA bins associated with ancestry after multivariate adjustment were evaluated for associations with mRNA abundance. Tumour purity was included in all mRNA models. Tumours with available mRNA abundance data were matched to those used in CNA analysis. For each gene affected by an ancestry-associated loss, its mRNA abundance was modeled against the ancestry of interest, copy number loss status, an ancestry-copy number loss interaction term, and tumour purity. The interaction term captures ancestry-associated mRNA changes. Statistical significance was assigned at q < 0.1. For genes affected by ancestry-associated gains, the same procedure was applied using gains. Complete mRNA modeling for CNAs is given in **Supplementary Table 5** and for SNVs in **Supplementary Tables 6**.

### Statistical Analysis & Data Visualization Software

All statistical analyses and data visualization were performed in the R statistical environment (v3.2.1) using the BPG^66^ (v5.9.8) package and with Inkscape (v0.92.3).

## Acknowledgments

The authors thank all the members of the Boutros lab for insightful discussions. This work was supported by the Discovery Frontiers: Advancing Big Data Science in Genomics Research program, which is jointly funded by the Natural Sciences and Engineering Research Council (NSERC) of Canada, the Canadian Institutes of Health Research (CIHR), Genome Canada and the Canada Foundation for Innovation (CFI). P.C.B. was supported by a Terry Fox Research Institute New Investigator Award and a CIHR New Investigator Award. This work was supported by an NSERC Discovery grant and by Canadian Institutes of Health Research, grant #SVB-145586, to PCB. The results described here are in part based upon data generated by the TCGA Research Network: http://cancergenome.nih.gov/. This work was supported by the NIH/NCI under award number P30CA016042 and by an operating grant from the National Cancer Institute Early Detection Research Network (1U01CA214194-01).

## Author Contributions

CHL and PCB initiated the project. CHL, and SH analyzed data. PCB supervised research. CHL and PCB wrote the first draft of the manuscript, which all authors edited and approved.

