## Supplementary Figures for "Ancestry Influences on the Molecular Presentation of Tumours"

### Supplementary Figures 1-4

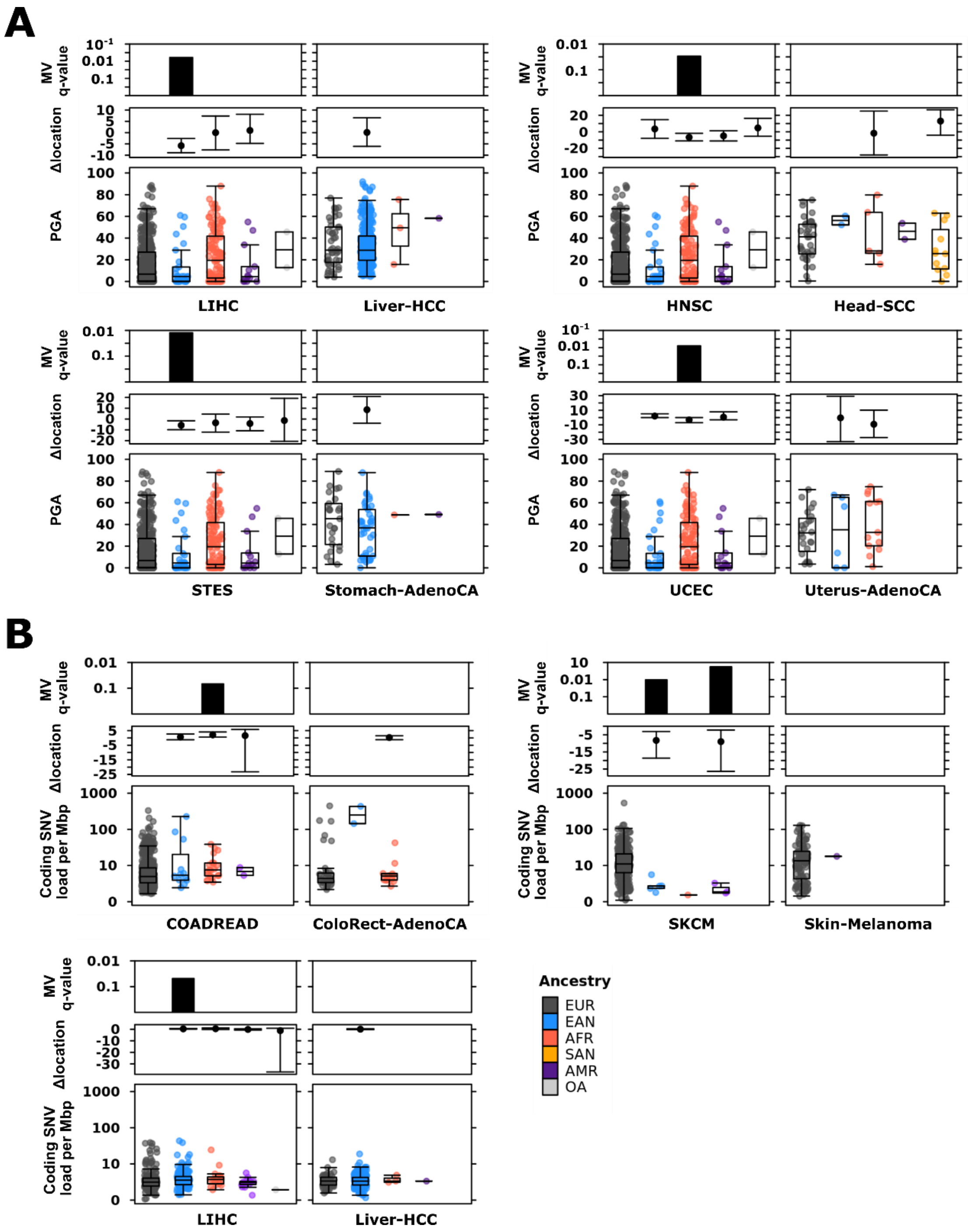


**Supplementary Figure 1. Ancestry-associated differences in mutation density.** Ancestry was associated with **(A)** percent genome altered and **(B)** SNV density in TCGA tumour-types, with corresponding PCAWG tumour-types for comparison. From top to bottom, each plot shows the adjusted multivariate p-value, difference in location, and mutation density. Tukey boxplots are shown with the box indicating quartiles and the whiskers drawn at the lowest and highest points within 1.5 interquartile range of the lower and upper quartiles, respectively.

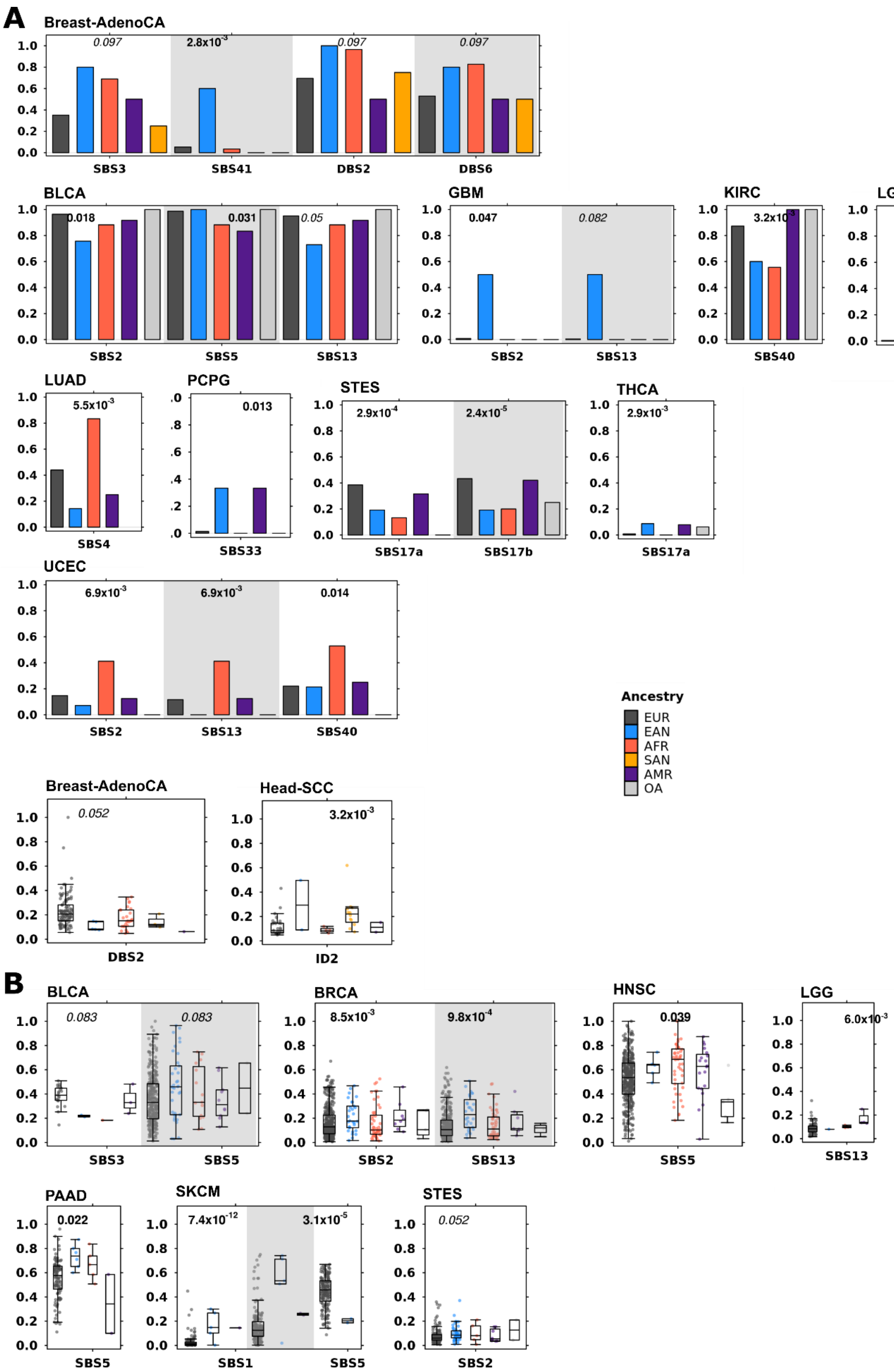

A
Breast-AdenoCA
0.097
2.8x10-3
0.097
0.097
SBS3
SBS41
DBS2
DBS6
BLCA
0.018
0.031
0.05
SBS2
SBS5
SBS13
GBM
0.047
0.082
SBS2
SBS13
KIRC
SBS40
LGG
0.029
SBS16
LUAD
5.5x10-3
SBS4
PCPG
0.013
SBS33
STES
2.9x10-4
2.4x10-5
SBS17a
SBS17b
THCA
2.9x10-3
SBS17a
UCEC
6.9x10-3
6.9x10-3
0.014
SBS2
SBS13
SBS40
Ancestry
EUR
EAN
AFR
SAN
AMR
OA
Breast-AdenoCA
0.052
DBS2
Head-SCC
3.2x10-3
ID2
B
BLCA
0.083
0.083
SBS3
SBS5
BRCA
8.5x10-3
9.8x10-4
SBS2
SBS13
HNSC
0.039
SBS5
LGG
6.0x10-3
SBS13
PAAD
0.022
SBS5
SKCM
7.4x10-12
3.1x10-5
SBS1
SBS5
STES
0.052
SBS2

**Supplementary Figure 2. Ancestry-associations with mutational signatures (A)** Ancestry-associations with signature detection. Barplots show frequency of signature detection in each ancestry group. **(B)** Ancestry-associations with signature relative activity. Tukey boxplots show relative signature activity as proportion of mutations attributed to each signature. Adjusted multivariate p-values shown above relevant ancestry group.

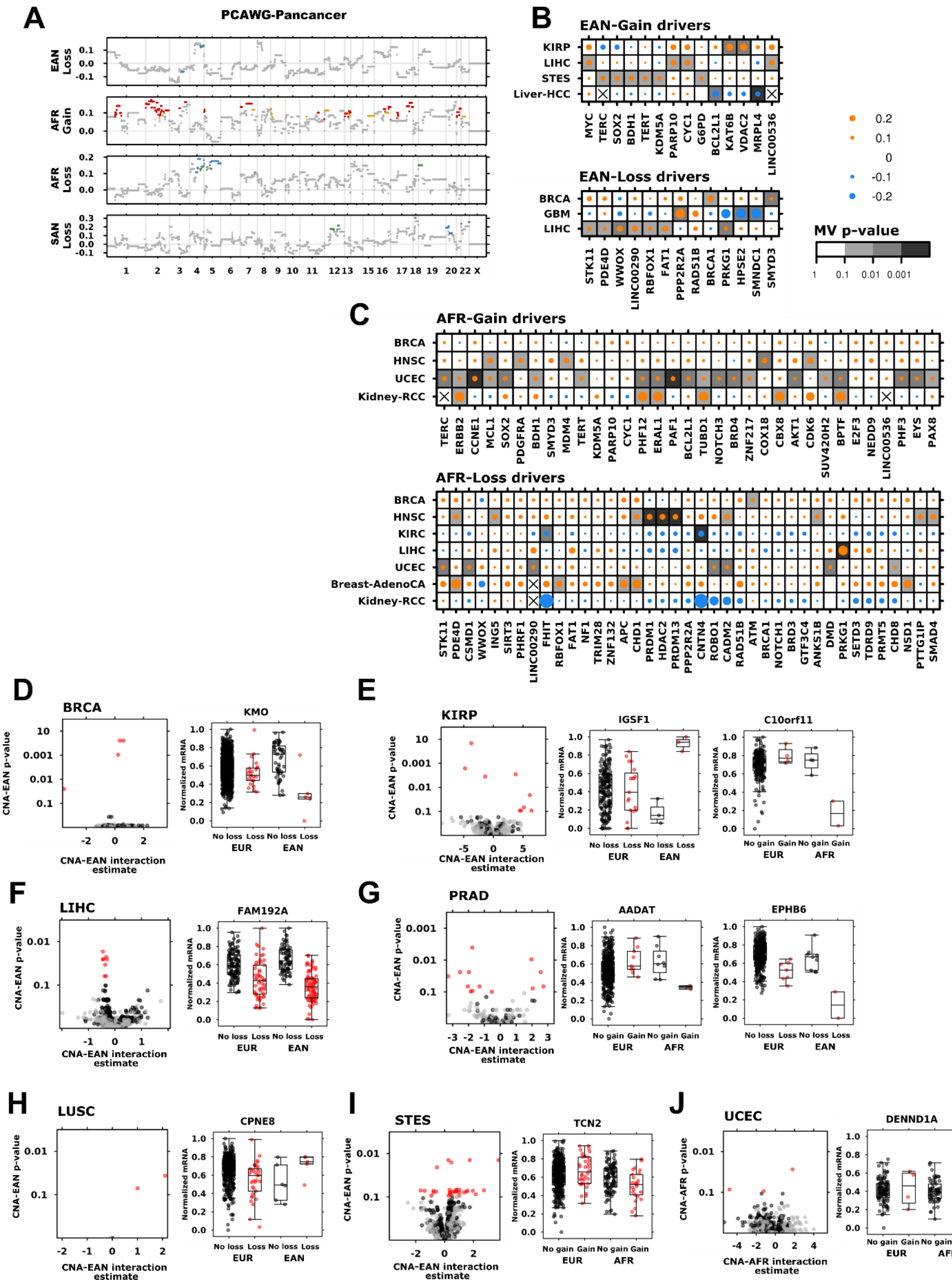

**Supplementary Figure 3. Ancestry-associations with CNAs and their downstream impacts on the transcriptome. (A)** Summary of ancestry associations in pan-PCAWG data. Each plot shoes the logistic regression coefficient estimate (for age analyses) or difference in proportion (for ancestry analyses) for the indicated variable and CNA type. Dot colour indicates statistical significance, where red (copy number gain) and blue (copy number loss) show adjusted $p < 0.05$ and yellow (gain) and green (loss) show adjusted $p < 0.1$. **(B)** EAN- and **(C)** AFR-associations with driver CNAs. **(D)**-**(I)** EAN- and **(J)** AFR-associated CNAs are themselves associated with changes in mRNA in each indicated tumour-type. For each tumour-type, the leftmost volcanoplot shows the adjusted p-value plotted against the coefficient of the CNA-age interaction for mRNA abundance, with each point representing a gene. Black dots show significant associations between mRNA and CNA; red dots show significant CNA-ancestry interactions. The right boxplots show mRNA abundances changes between CNA (red) or no CNA (loss) in tumours of **(D)**-**(I)** EUR and EAN or **(J)** EUR and AFR ancestry. Tukey boxplots are depicted.

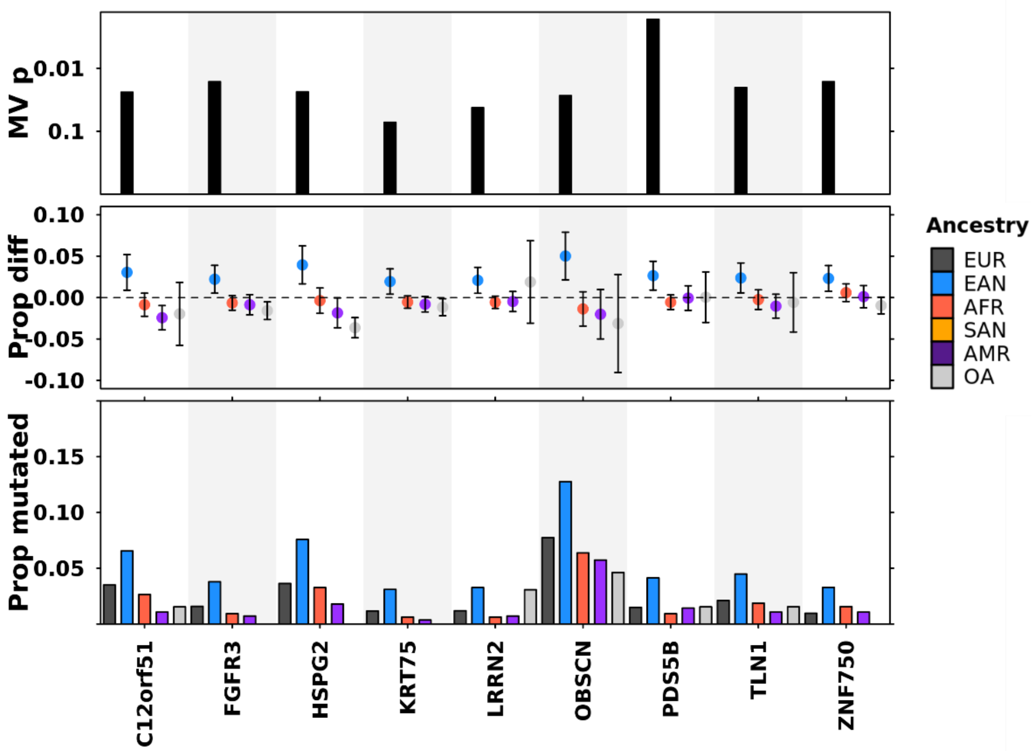


**Supplementary Figure 4. Associations between host factors and SNVs in pan-TCGA analysis.** Pan-TCGA associations between ancestry and SNV frequency. Genes with ancestry - biased mutation frequencies shown with top showing adjusted multivariate p-values, middle showing difference in proportion, and bottom showing proportion of tumours with mutated gene per ancestry.
